# Mycelium-bound lipases of psychrophilic and mesophilic fungi as biocatalysts for biodiesel synthesis

**DOI:** 10.1101/2023.11.28.568820

**Authors:** Mateusz Kutyła, Natalia Jaszek, Wiktoria Jędrys, Sandra Graba, Ewelina Pluta, Katarzyna Gdula, Aleksandra Batyra, Amelia Szczepańska, Alicja Śliwa, Laura Cieślak, Anna Marzec-Grządziel, Mariusz Trytek

## Abstract

In this study, biocatalysts in the form of mycelium-bound lipases of psychrotrophic and mesophilic fungi were screened for their ability to synthesize fatty acid methyl (FAME) and ethyl esters (FAEE). It has been shown that the hydrolytic activity of biocatalysts is not accompanied by their biodiesel synthetic activity. Preliminary studies showed for the first time, that the FTIR technique can be used to predict synthetic activity. The highest correlation between the intensity of the bands and synthetic activity, negative and positive, respectively, was observed for two neighboring bands at 1745 cm^-1^ (*R* = −0.73) and 1699 cm^-1^ (*R* = 0.68). The psychrotrophic strain *Penicillium* sp. 59, isolated from the soil of the Arctic tundra (Spitsbergen), has been shown for the first time as an efficient whole-cell biocatalyst for fatty acid ethyl ester synthesis and is stable in organic solvents. Mathematical optimization of biosynthesis was also performed using RSM. A molar canola oil conversion value of 83.3% into FAEE was obtained at 24.2 °C in n-hexane using 12.5% of used canola oil and 5.3% of the fungal biocatalyst within 6 h. Dry mycelium of *Penicillium* sp. 59 is a promising new biocatalyst for large-scale biosynthesis of biodiesel, given its low-cost production, high activity at room temperature, no need to mix reactions, and reusability in a minimum of six cycles.

## 1. INTRODUCTION

Demographic growth has resulted in an increase in worldwide energy demand. According to the U.S. Energy Information Administration (EIA), the estimated energy consumption between 2015 and 2040 will increase by 28%, and by 2050, nonrenewable fossil fuels will supply 65% of the world’s energy needs [1,2]. In the Americas, Europe, and Africa, oil remains the main source of energy for transportation and industry [2]. Burning fossil fuels contributes to climate change and environmental pollution by emitting greenhouse gases and other harmful pollutants, such as NO_x_, SO_x_ and CO [3]. Biofuels, including biodiesel, are viable alternatives to the conventional fossil fuels. Increasing the use of renewable energy sources in the form of biodiesel would not only help address the global energy crisis and reduce pollution, but would also help to manage industrial and household waste, such as used cooking oil, which can provide the feedstock for this biofuel [3].

Biodiesel is a methyl (FAME) or ethyl fatty acid ester (FAEE) obtained by the transesterification of triglycerides and alcohol [1]. Therefore, suitable catalysts are required for biodiesel synthesis. Heterogeneous catalysts have high selectivity and activity, can be used repeatedly, are easy to store, and exhibit low toxicity. Homogeneous catalysts are divided into alkaline (NaOH, KOH) and acidic (H_2_SO_4_, H_3_PO_4_) catalysts, which are dissolved in the reaction mixture, making their reuse difficult [4]. Homogenous processes are more expensive than heterogeneous ones. The chemical synthesis of biodiesel has many disadvantages, including high water and energy consumption, and environmental pollution [5]. Therefore, attempts have been made to replace chemical methods with enzymatic processes, which are safer for the environment. The biocatalyst used in biodiesel synthesis should satisfy the following conditions: mechanical and thermal stability, non-stereospecificity, reusability, and low cost [6].

Lipases (EC 3.1.1.3), which are mainly derived from filamentous fungi, yeasts, and bacteria, are used as biocatalysts for biodiesel synthesis. These enzymes are stable under harmful chemical and physical conditions [7]. Currently, many lipases from filamentous fungi, such as *Rhizomucor miehei* (RML), *Thermomyces lanuginosus* (TLL), *Rhizopus oryzae* (ROL), *Aspergillus niger* (ANL), and *Penicillum expansum* lipase (PEL), are used as biocatalysts in research and biodiesel production. Many of these are commercially available, usually as immobilized enzymes [8]. In non-aqueous media, lipases catalyze esterification, acidolysis, aminolysis, and interesterification reactions with high selectivity and specificity. Biocatalysis typically occurs under mild conditions (temperature 20-60 °C). The use of lipases in biodiesel synthesis allows for a simplified production process, easy purification of the main product, and isolation of glycerol by-products. Because lipases can catalyze esterification and transesterification reactions, it is possible to produce high-quality biodiesel from inexpensive feedstock [8].

Fungal lipases have many advantages, but their use has some limitations. The cost of obtaining pure enzymes is high. Efficient biocatalysis requires the use of immobilized enzymes to increase the stability and reusability of biocatalysts [8]. However, immobilization is associated with significant financial costs and the risk of enzyme activity loss during the process. To minimize these drawbacks, it is possible to use a biocatalyst in the form of mycelium-bound lipase for the synthesis of biodiesel [9–11]. Thus, there is no need for isolation, purification, or additional immobilization of enzymes on the selected carriers. Biocatalysts of this type are easy to obtain and inexpensive. Mycelium-bound lipases are characterized by greater stability because the reaction is carried out in an environment close to natural conditions, ensuring that the enzymes are protected from negative external influences. In addition, the reaction carried out using this type of biocatalyst facilitates several reaction cycles, owing to its ease of recovery [8,12,13].

Despite the availability of a large number of commercial lipases, there is still a need to search for new lipase sources, especially psychrophilic microorganisms. The use of psychrophilic lipases reduces the process temperature, which reduces the energy consumption and carbon footprint during biodiesel production. Unfortunately, screening of new biocatalysts in the form of whole cells is time-consuming and makes it impossible to test a large number of strains in a short time.

In this paper, different psychrophilic and mesophilic fungal species as biocatalysts for the biosynthesis of biodiesel were studied. Herein, we propose for the first time the use of FTIR-ATR analysis of the mycelium to improve or shorten the screening of biocatalysts by correlating the intensity of the bands with the biosynthetic activity of the mycelium. The synthesis of fatty acid ethyl esters by the novel psychrotrophic fungus *Penicillium* spp. 59 was mathematically optimized.

## 2. MATERIALS AND METHODS

### 2.1. Chemicals

Tributyrin (97%), calcium carbonate (≥99%), and Tween 85 (99%) were purchased from Sigma, USA). K_2_HPO_4_, MgSO_4_ × 7H_2_O, yeast extract, (NH_4_)_2_SO_4_, *n*-hexane, chloroform, ethanol, heptane, isopropanol, methanol, ethyl acetate, and toluene (≥99% purity) were obtained from POCH, Poland. Olive oil was sourced from a local grocery store. Used canola oil was obtained from a local restaurant.

### 2.2. Microorganisms and media

Mesophilic *Aspergillus oryzae* CBS 133.52, *Trichoderma harzianum* CBS 688.94, *T. harzianum* CBS 415.68, *T. harzianum* CBS122771, *T. harzianum* CBS 416.96, *T. harzianum* CBS 436.95, *T. harzianum* CBS 123057, *Neurospora crassa* 4/4, *A. wenti* /1, *Rhizopus oryzae* DSM 63539, *R. oryzae* NRLL 3613, *Rhizomucor miechei*, *T, reesei* F560, *Fusarium solani v. radicule* and strains of psychrophilic fungi isolated from Arctic tundra (West Spitsbergen) soil were labeled as 14, 18, 24, 25, 36, 37, 46, 49, 51, 59, 65, 66, 75, 77, and 96. Prior to the experiments, fungi had been maintained on agar slants at 4 °C and subcultured every month.

Fungi were cultivated on the basal medium (BM) composed of 0.7% Tween 85, 0.1% yeast extract, 0.5% (NH_4_)_2_SO_4_, 0.1% K_2_HPO_4_, 0.02% MgSO_4_ × 7H_2_O, 1% olive oil, and 0.5% CaCO_3_ (pH = 6.0, adjusted by adding 0.1 M HCl) [13].

### 2.3. Submerged culture conditions

For the cultivation of fungi, 100 mL of sterile basal medium (in 300-mL Erlenmeyer flasks) was uniformly inoculated with 4 mL of fungal spore suspension (at a concentration of 4 × 10^5^ mL^-1^) and cultivated on a rotary shaker (150 rpm) for 4 days at 20 °C and 30 °C for psychrotrophic and mesophilic fungi, respectively [13].

After completion of the culture, the mycelia were separated from the post-culture liquid by centrifugation at 10,000 *g* for 15 min at 4 °C. The supernatant was used as the source of extracellular lipase. The mycelial biomass was frozen at –20 °C overnight and then lyophilized for 24 h. Freeze-dried mycelia were milled and used as intracellular lipases. Mycelium-bound lipase has also been used for the biocatalytic synthesis of biodiesel and for the FTIR-ATR analysis.

### 2.4. Determination of extracellular and intracellular lipolytic activity

Extracellular lipase activity was determined according to a previously described method [13]. Briefly, the supernatant (0.5 mL) was mixed with 1 mL of 50 mM phosphate buffer (pH 7.0) and olive oil (1.5 mL) in a 25-mL Erlenmeyer flask. The samples were shaken (200 rpm) for 1 h at 30 °C, and then 1 mL of 96% ethanol was added. The concentration of the free fatty acids was determined by titration with 0.0500 M KOH. The control was a reaction with heat-inactivated enzyme.

Intracellular lipolytic activity was determined according to the procedure described above, with the exception that 10 mg of lyophilized mycelium was used as the enzyme source instead of the supernatant. One unit of lipase activity (U) was defined as the amount of enzyme that released 1 *μ*mol · min^-1^ of free fatty acids. Extracellular and intracellular lipase activity were expressed as U/mL of the post-culture and U/g of fungal lyophilizate, respectively.

### 2.5. Screening of biocatalysts towards the synthesis of methyl (FAME) and ethyl (FAEE) esters

The biocatalytic synthesis of biodiesel was carried out in sealed 25-mL Erlenmeyer flasks equipped with magnetic stirring bars. In the standard procedure, 173 *μ*L of used canola oil and 30 *μ*L of ethanol or 25 *μ*L of methanol were dissolved in 2.5 mL of hexane containing 2% (w/v) of the fungal biocatalyst and incubated at 40 °C for 24 h.

### 2.6. Analysis of the biodiesel synthesis efficiency

After 24 h of biocatalytic synthesis, 100 *μ*L of the reaction mixture was collected, dried over anhydrous sodium sulfate, and diluted tenfold with n-hexane containing tetradecane as an internal standard at a concentration of 1 g/L. The samples were then subjected to GC-FID and GC-MS analyses. The analyses were conducted using DB−5 capillary column (30 m × 0.25 mm inner diameter). The separation conditions were as follows: helium was used as the carrier gas at a constant flow rate of 1 mL/min and the injection temperature was 290 °C. The oven temperature program was as follows: 180°C for 2 min, which was then increased at a rate of 5 °C/min to 230 °C for 5 min, and then increased at a rate of 10 °C/min to 290 °C for 5 min. The biodiesel synthesis efficiency was expressed as the molar conversion (%) of used canola oil into fatty acid esters.

### 2.7. Fourier-transform infrared spectroscopy

The dry mycelia were analyzed using attenuated total reflection infrared spectroscopy (FTIR-ATR) to examine the qualitative and quantitative differences between biocatalyst spectra. The spectra were recorded using a TENSOR 27 Bruker spectrometer equipped with a diamond onto which the tested mycelium was applied. The spectra were recorded in the wavenumber range 4000 – 600 cm^-1^ with 60 scans per spectrum at a resolution of 1 cm^-1^.

### 2.8. Phylogenetic analysis of the psychrotrophic fungus strain 59

DNA from fungal culture was isolated with commercially available kit (Bead-Beat Micro AX Gravity, A&A Biotechnology, Poland). Purity and concentration of isolated DNA were checked with fluorimeter (Quantus™ Fluorometer, Promega, Germany). PCR conditions for ITS region amplification were as follows: 95 °C – 15 min; 30 cycles for 95 °C – 30 sec, 50 °C – 1 min, 72 °C – 1.5 min; 72 °C – 7 min. Mix reagents were: 4 µL Silver Hot Start PCR MIX LOAD (Syngen Biotech, Poland), ITS1 primer 0.25 µM, ITS4 primer 0.25 µM, 14 µL H2O, 1 µL DNA. PCR product was checked on agarose gel (1.5%, 100 V, 1 h) for purity. Sanger sequencing was prepared as an external service (Genomed, Poland). Fasta sequences for both primers were analyzed using BioNumerics software (Applied Maths N.V., Belgium). Merged sequence was aligned into database (NCBI) using Seed2 [14].

### 2.9. Optimization of biodiesel synthesis

In the first step of optimization, the influence of the type of solvent was determined on the efficiency of biodiesel synthesis by strains of the fungus 59 and *Aspergillus oryzae*.

In the optimization experiments, independent variables were placed at one of three spaced values coded as −1, 0, and 1, and response surface methodology (RSM) using R programming language was used. A six-factor [volume of canola oil (*X*1), volume of ethanol (*X*2), biocatalyst amount (*X*3), temperature (*X*4), rotation of the reaction (*X*5), and volume of the solvent (*X*6)] and mixed-level [*X*1 (90, 180, and 270 *μ*L), *X*2 (30, 60, and 90 *μ*L), *X*3 (25, 75, and 125 mg), *X*4 (20, 40, and 60 °C), *X*5 (0, 50, and 100 RPM), *X*6 (1.5, 2.5, and 3.5 mL)] design were used in the experiments (Table 1). In total, 53 biodiesel synthesis reactions were carried out to obtain a response and determine the surface for the reaction yield. The reaction was performed for 6 h.

**Table 1.**
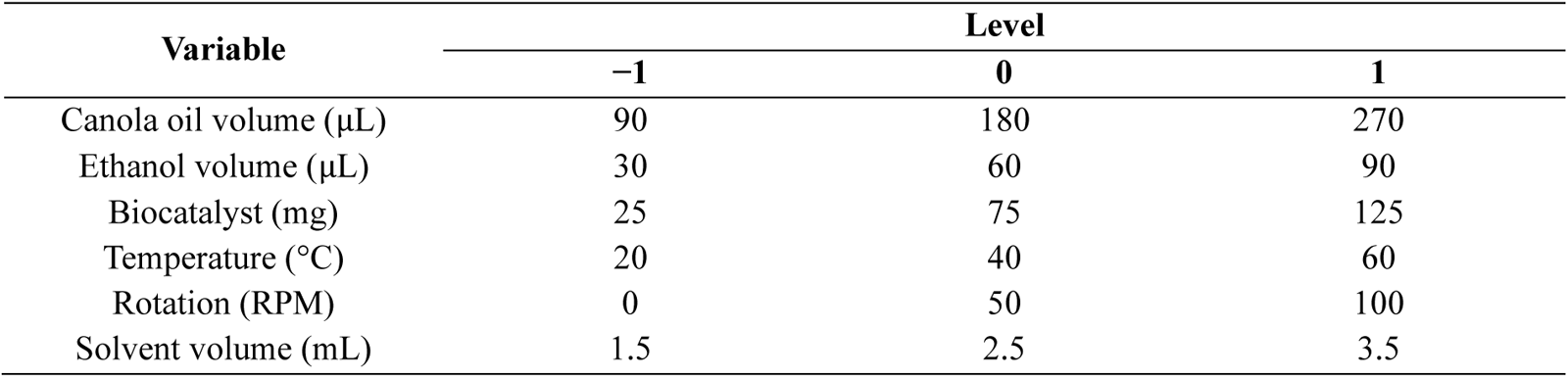
Independent variables and their level used in the design of the experiment.

| Variable | Level |  |  |
| --- | --- | --- | --- |
|  | -1 | 0 | 1 |
| Canola oil volume (μL) | 90 | 180 | 270 |
| Ethanol volume (μL) | 30 | 60 | 90 |
| Biocatalyst (mg) | 25 | 75 | 125 |
| Temperature (°C) | 20 | 40 | 60 |
| Rotation (RPM) | 0 | 50 | 100 |
| Solvent volume (mL) | 1.5 | 2.5 | 3.5 |

Based on empirical data, “FO” (first order), “TWI” (two-way interactions), “SO” (second order) and “PQ” (pure quadratic) models were created using R programming language and the “*rsm*” package [15].

### 2.10. Reusability of the fungal biocatalyst in the reaction of biodiesel synthesis

The mycelia of the fungus 59 strain (108 mg) were weighed into 25-mL glass flasks. Then an optimized amount of hexane (2.05 mL), canola oil (255.6 *μ*L) and ethanol (35.1 *μ*L) were added. The flasks were sealed with a silicone stopper and incubated for 6 hours at 24.2 °C. Then, 100 *μ*L of the reaction mixture was added to 900 *μ*L of hexane and analyzed using GC-FID. The mycelium was then filtered, rinsed twice with hexane, and used in the next catalytic cycle.

### 2.11. Statistical analysis

The experiments were performed in triplicate. The results were subjected to analysis of variance (ANOVA) followed by Tukey’s post-hoc test for multiple comparisons. The correlation between biodiesel synthesis activity and band intensity in the FTIR-ATR spectrum was analyzed using Pearson’s correlation coefficient *R*. Statistical analysis was performed using the R programming language version 3.6.1.

## 3. RESULTS

In the first stage of the study, fungi were cultured in a liquid medium containing Tween 85 and olive oil as lipase inducers. Extra- and intracellular lipolytic activity, protein concentration, specific enzyme activity, pH of the post-culture liquid, and mycelium dry weight were determined after the end of the culture. All tested fungi were able to grow on medium containing Tween 85 and olive oil. The amount of lyophilized mycelia obtained ranged from 4.5 to 19.1 g/L (Tab. 2). The highest extracellular lipolytic activity was shown by four strains: mesophilic *T. harzianum* CBS 436.95 (10.8 ± 1.8 U/mL) and psychrotrophic 66 (9.9 ± 1.2 U/mL), 96 (8.6 ± 0.6 U/mL) and 59 (8.3 ± 0.9 U/mL). Significant extracellular activity was also shown by five mesophilic strains of *F. solani v. radicule*: *R. michei*, *R. oryzae* DSM 63539, *R. oryzae* NRLL 3613, *T. harzianum* CBS 688.94, and psychrotrophic 77 (in the range of 3.3–4.7 U/mL). The other strains showed activity lower than 2.5 U/mL. The intracellular activity of fungal biomass was the highest in psychrotrophic strains 75, 59, 18, 14, 46 and 36 (ranging from 40.7 to 60.3 U/g), and two strains of *R. oryzae* (41 and 47 U/g) stood out among the mesotrophic strains. Acidification of the medium during cultivation was observed in nine of the tested strains, in which significant extracellular lipolytic activity was observed. Psychrotrophic strains are characterized by a higher synthesis of extracellular proteins compared to the mesophilic strains studied. Only the protein concentration in the culture fluid of strain 59 was 9.4 µg/mL, which was accompanied by the highest specific activity (Table 2).

**Table 2.**
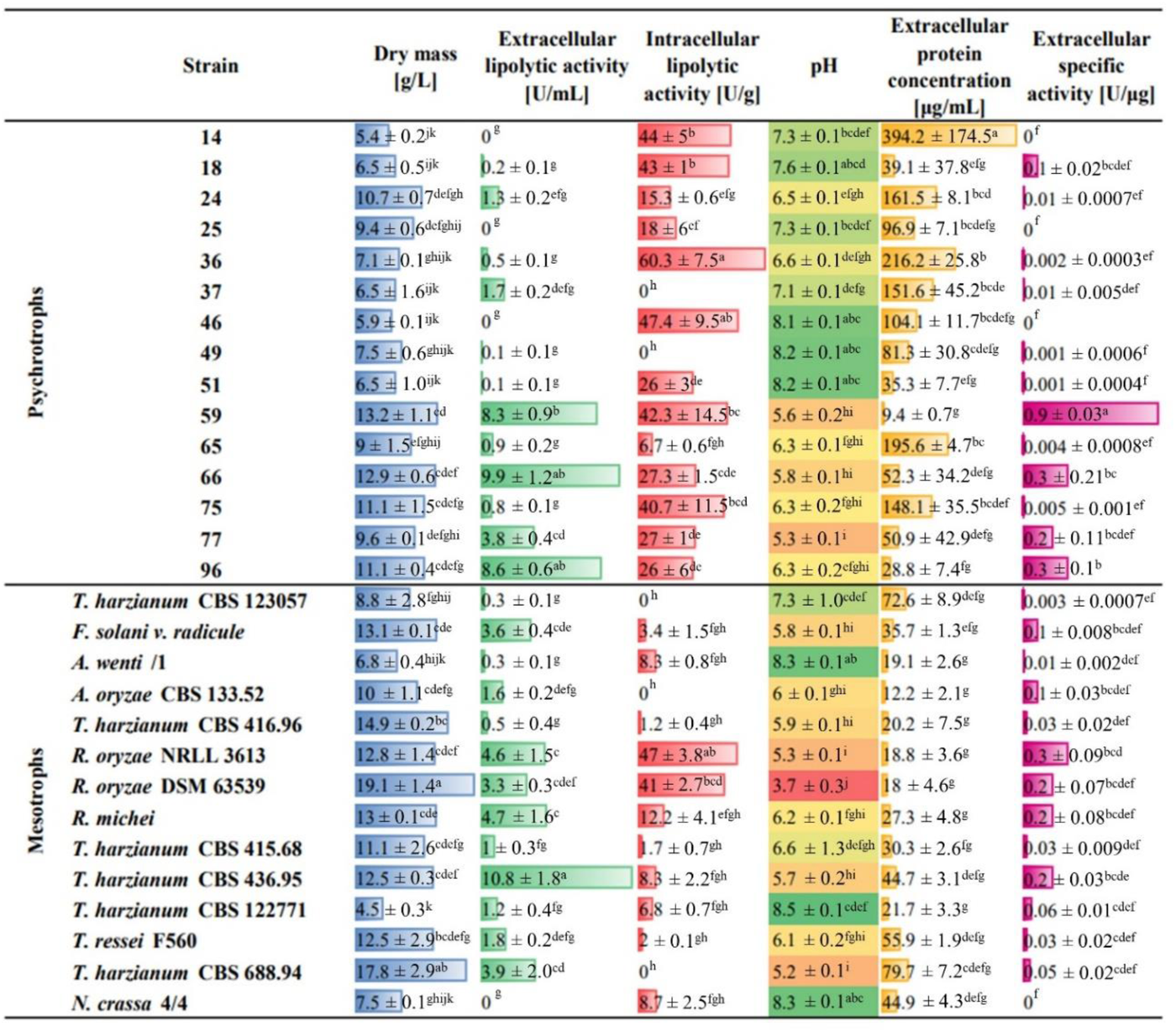
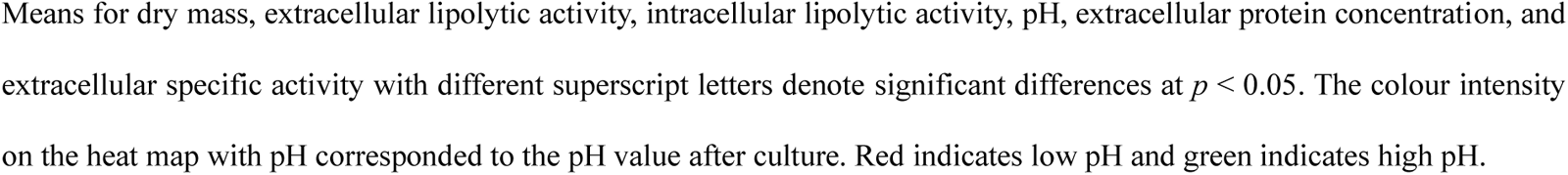
Dry mass, extra- and intracellular lipolytic activity, pH, protein concentration and specific activity obtained after cultivation of psychrotrophic and mesophilic filamentous fungi.

In the next stage of research, freeze-dried mycelia were used as biocatalysts in the synthesis of fatty acid methyl (FAME) and ethyl (FAEE) esters. Only 4 out of 29 strains showed significant activity in biodiesel synthesis, and these were psychrotrophic 59 (57.5 ± 3.6% FAME and 90.1 ± 7.3% FAEE yield), and mesophilic *A. oryzae* CBS 133.52 (91.1 ± 8.3% FAME and 94.5 ± 1.5% FAEE yield), *R. oryzae* DSM 63539 (75.8 ± 1.6% FAME and 76.8 ± 3.9% FAEE yield) and *R. oryzae* NRLL 3613 (75.8 ± 0.6% FAME and 79.3 ± 2.4% FAEE yield) (Fig. 1). The mesophilic strains showed similar methyl and ethyl ester synthesis activities. Among the psychrotrophic strains, approximately 45% of the fatty acid ethyl ester synthesis activity was also shown by 66 (43.4 ± 3.3%) and 96 (48.3 ± 1.8%).

**Figure 1.**
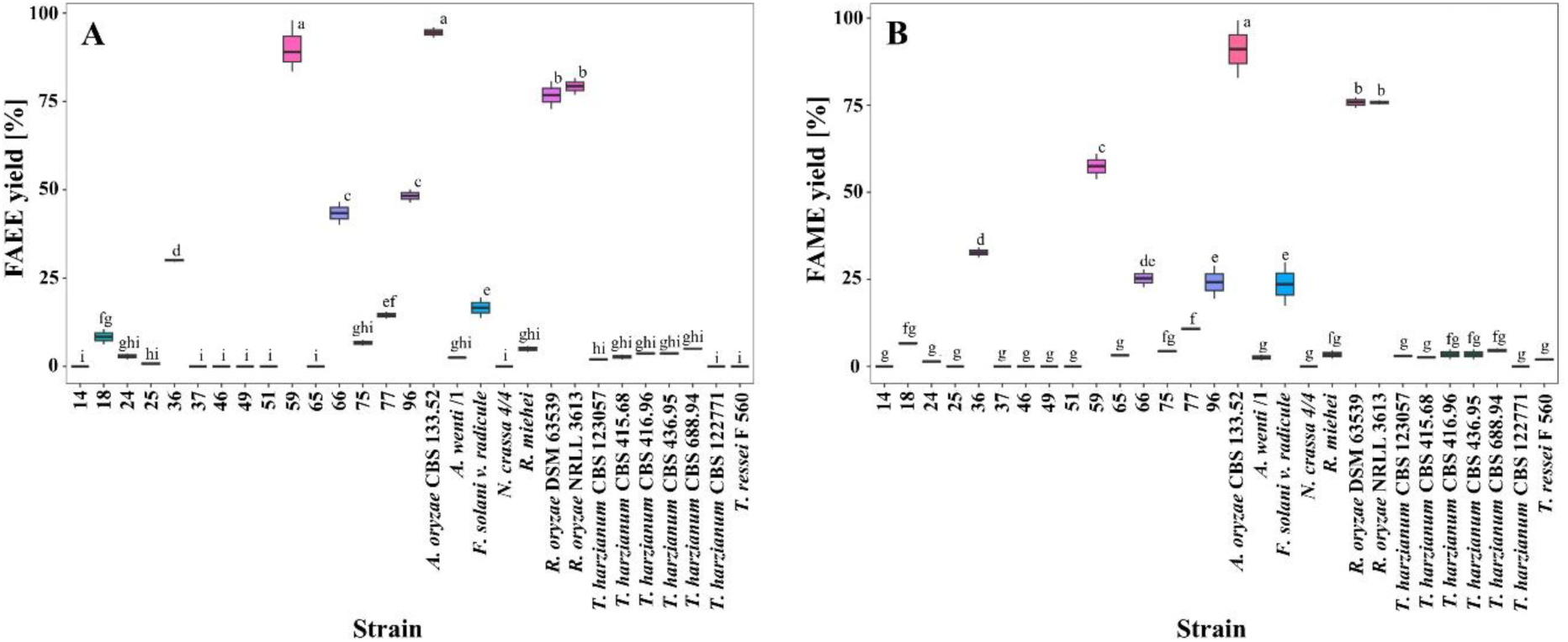
Efficiency of the synthesis of ethyl (A), and methyl (B) fatty acid esters by mycelia of psychrotrophic and mesophilic fungi. Means for FAEE and FAME yields with different superscript letters denote a significant difference at *p* < 0.05

To fully illustrate the obtained data and find the variable most closely related to the synthetic activity of mycelia, a correlation matrix was created from the results obtained for all strains. The increase in the biomass obtained in one culture cycle was strongly negatively correlated (*R* = −0.85) with the pH of the postculture fluid. A negative correlation *R* ranging from −0.57 to −0.55 was also obtained between the pH of the culture fluid and the synthetic activity of methyl and ethyl esters. The activity of the fungal biocatalyst increased with decreasing pH of the postculture fluid. Importantly, a low positive correlation was found between the extra- and intracellular lipolytic activity and the synthetic activity of mycelia. The high hydrolytic activity of the mycelia did not translate into their equally high synthetic activity. In contrast, intracellular hydrolytic activity was not strongly correlated with any variable (Fig. 2). Due to the lack of clear relationships between synthetic activity and other data obtained after cultivation, in the next experiment, we determined the FTIR spectrum of fungal biocatalysts and examined whether it was possible to isolate a band (or bands) strongly correlated with synthetic activity. Such a correlation would allow for the use of an inexpensive and fast method to test mycelia for use as an efficient biocatalyst for biodiesel synthesis.

**Figure 2.**
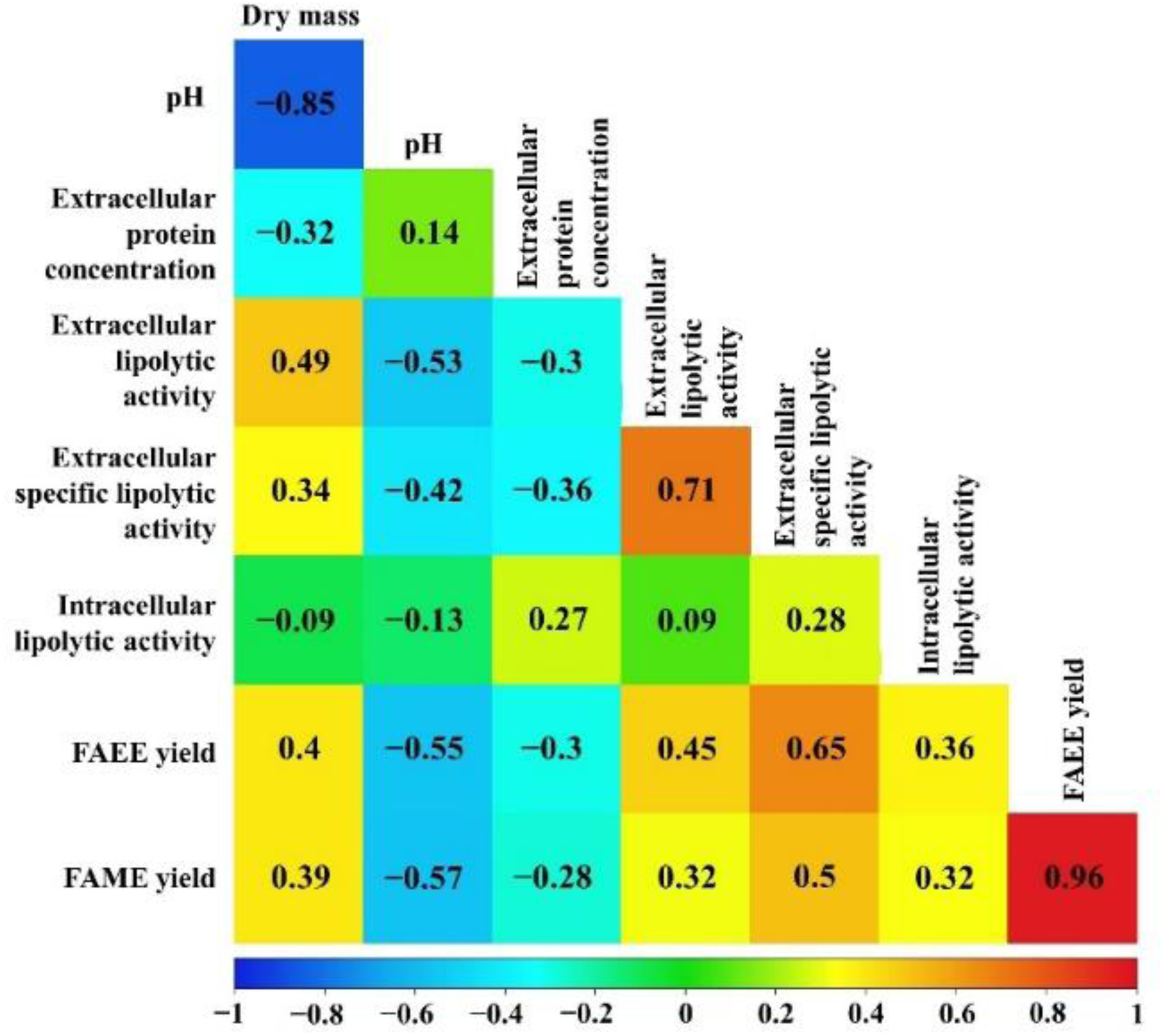
Correlation matrix between pH, extracellular protein concentration, extracellular lipolytic activity, extracellular specific lipolytic activity, intracellular lipolytic activity (obtained in cultures of all psychrotrophic and mesophilic fungi), and FAEE yield and FAME yield. The results were presented as Pearson’s correlation coefficient *R*

Analysis of the correlation between the intensity of the bands in the FTIR spectrum of the fungal biocatalysts and their activity in biodiesel synthesis indicated bands that had a significant impact on the activity of the mycelium. A high correlation (*R* > 0.6, *R* < −0.6, *p* > 0.001) was found between the bands at 2690–2530, 1745, 1699, 1161, and 790 cm^-1^ and the synthetic activity of the biocatalyst. The highest negative correlation (*R* = −0.73) was recorded for the 1745 cm^-1^ band and the highest positive correlation (*R* = 0.68) for the adjacent 1699 cm^-1^ band (Fig. 3). The bands at 1745 and 1699 cm^-1^ are characteristic of the stretching vibrations of the C=O groups. The bands at 2680–2530 cm^-1^ indicate the presence of O–H or N–H groups. The band at 1161 cm^-1^ occurs in the presence of C–O or C–N chemical groups. In turn, the N–H or O–N groups may indicate the presence of a band at 790 cm^-1^ [16].

**Figure 3.**
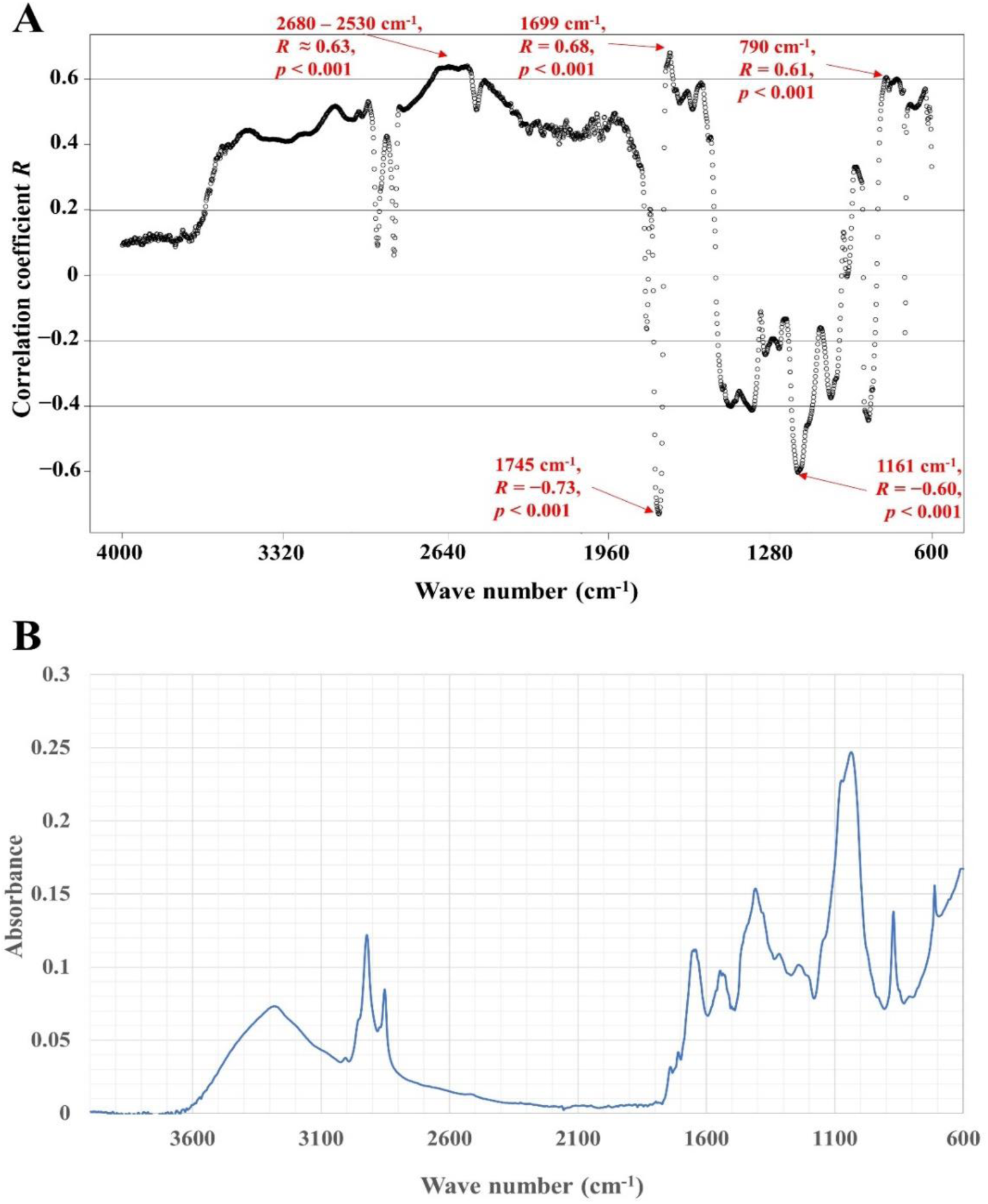
Graph of the distribution of the correlation coefficient *R* depending on the wave number and synthetic activity of biodiesel (A), and FTIR-ATR spectra of dry mycelium from psychrotrophic strain 59 (B). All 29 fungal strains were analyzed in at least three replicates.

Because of the high synthetic activity of biodiesel of the new cold-adapted fungus 59, we decided to conduct its phylogenetic analysis. Based on ITS sequence and database comparison, strain 59 belongs to the genus *Penicillium*. Table 3 provides information on the best matches for the given species.

**Table 3.** Similarity and coverage of the ITS sequence of the psychrotrophic strain 59 to the sequence of *Penicillium* species.

| Accession | Species | Similarity [%] | Coverage [%] | Query length |
| --- | --- | --- | --- | --- |
| KP411588 | *Penicillium echinulatum* T3.10-3 | 99.1 | 86.7 | 788 |
| KP411569 | *Penicillium crustosum* C2-5.10.1.S2 | 98.7 | 86.5 | 788 |
| KP411582 | *Penicillium commune* T.S1.F | 98.4 | 86.9 | 788 |
| KP857656 | *Penicillium* sp. AQG11 | 98.8 | 84.1 | 788 |
| KU216705 | *Penicillium rubens* XQ3 | 97.8 | 87.1 | 788 |
| KU216703 | *Penicillium rubens* XQ1 | 98.1 | 86.2 | 788 |
| MF034654 | *Penicillium griseofulvum* L1 | 98.5 | 84.9 | 788 |
| KU216725 | *Penicillium chrysogenum* XQ23 | 98 | 86.8 | 788 |
| KF597019 | *Penicillium polonicum* NFW9 | 97.8 | 86.4 | 788 |
| KX421790 | *Penicillium polonicum* TLN1 | 98.1 | 85.7 | 788 |

As the next step in this study, the effect of organic solvents on the synthetic activity of ethyl esters of two types of fungi, *Penicillium* sp. 59 and *Aspergillus oryzae*, was analyzed. The best solvents for biodiesel synthesis for both species were hexane and heptane. Transesterification occurred in approximately 50% yield in toluene. The use of ethyl acetate to induce interesterification resulted in approximately 25% conversion of the oil to fatty acid ethyl esters. The biocatalyst in the form of *A. oryzae* mycelia showed trace activity in chloroform and isopropanol but not in ethanol and methanol. Importantly, the *Penicillium* sp. 59 mycelia showed approximately 30% activity in ethanol. The addition of ethanol to ethyl acetate as a solvent results in an approximately 60% synthesis yield. No catalytic activity was observed for methanol (Fig. 4).

**Figure 4.**
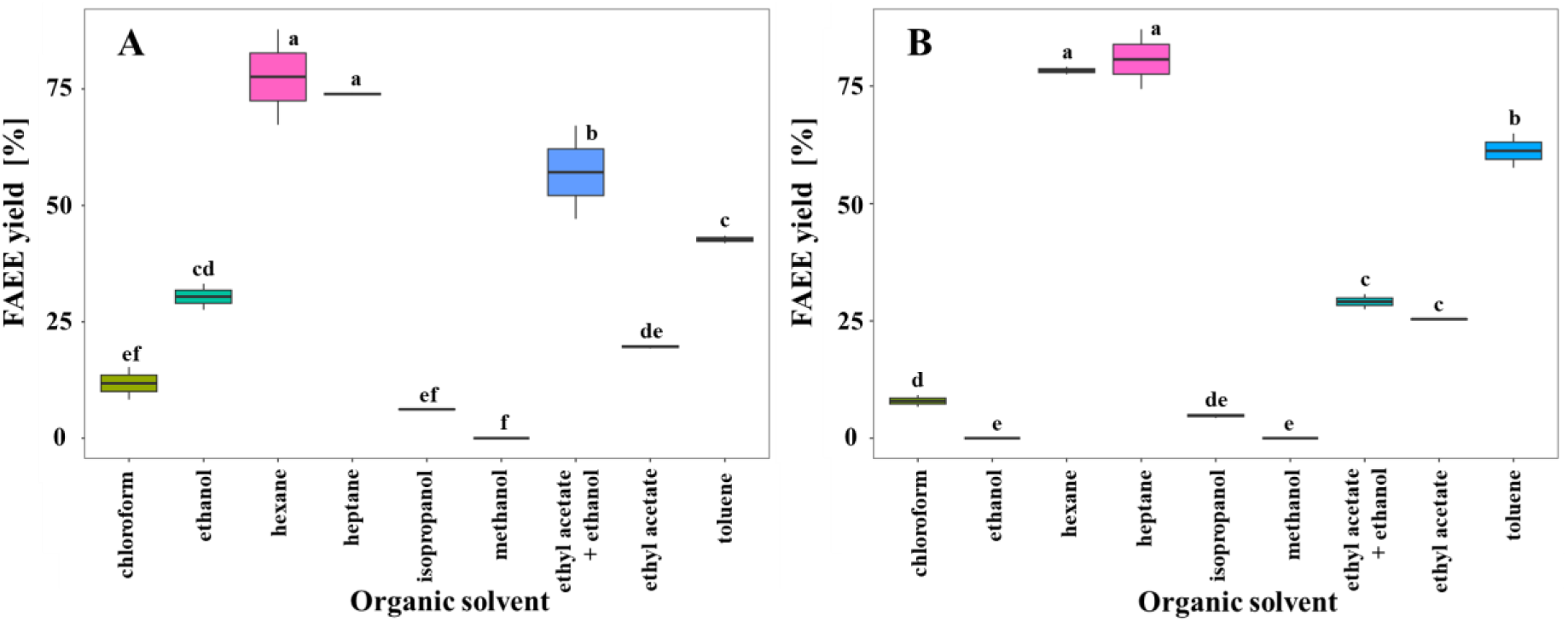
Influence of the type of solvent on biodiesel synthesis efficiency by mycelium of psychrotrophic *Penicillium* sp. 59 (A) and mesotrophic *Aspergillus oryzae* CBS 133.52 (B)

Based on the empirical results, four generated models were developed for optimization of biodiesel synthesis and experimentally tested: first-order, two-way interaction, second-order, and pure quadratic. The models used are listed in Table 4. In all models, neither the volume of solvent in the range of 30– 90 *μ*L nor the mixing of the system had significant effect (*p* < 0.05) on the efficiency of biodiesel synthesis. The second-order model was characterized by the highest coefficient of determination (*R*^2^ = 0.92), while the prediction based on this model significantly differed from the experimental data, which is why it was rejected. A two-way interaction model was selected as the best model for the synthesis of fatty acid ethyl esters by *Penicillium* sp. 59. The real FAEE yield was the highest among the four models and was close to the predicted yield in this model. The two-way interaction model coefficients are presented in Table 5.

**Table 4.**
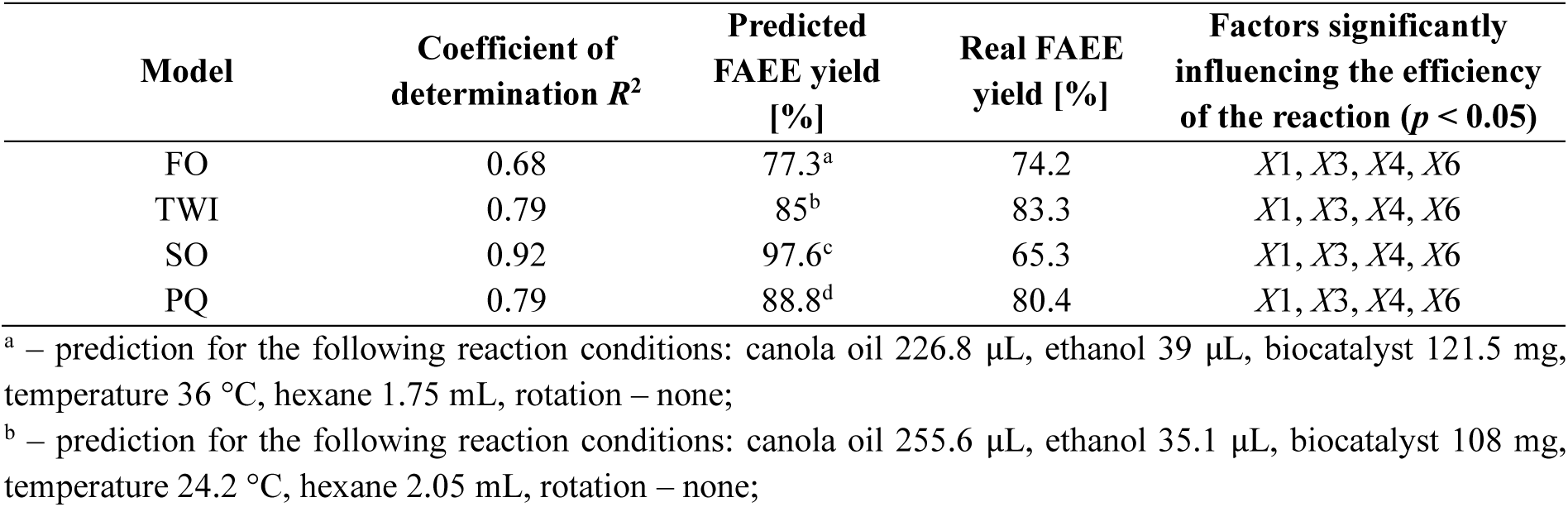
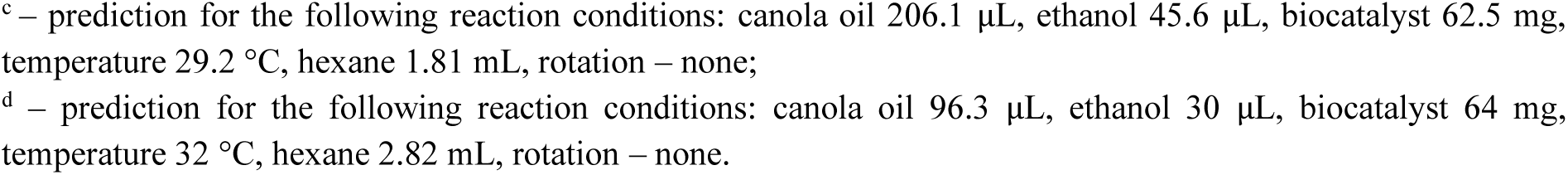
Optimization models for prediction of FAEE yield.

| Model | Coefficient of determination $R^2$ | Predicted FAEE yield [%] | Real FAEE yield [%] | Factors significantly influencing the efficiency of the reaction ($p < 0.05$) |
| --- | --- | --- | --- | --- |
| FO | 0.68 | 77.3<sup>a</sup> | 74.2 | $X_1, X_3, X_4, X_6$ |
| TWI | 0.79 | 85<sup>b</sup> | 83.3 | $X_1, X_3, X_4, X_6$ |
| SO | 0.92 | 97.6<sup>c</sup> | 65.3 | $X_1, X_3, X_4, X_6$ |
| PQ | 0.79 | 88.8<sup>d</sup> | 80.4 | $X_1, X_3, X_4, X_6$ |

<sup>a</sup> – prediction for the following reaction conditions: canola oil 226.8 $\mu\text{L}$ , ethanol 39 $\mu\text{L}$ , biocatalyst 121.5 mg, temperature 36 $^{\circ}\text{C}$ , hexane 1.75 mL, rotation – none;
<sup>b</sup> – prediction for the following reaction conditions: canola oil 255.6 $\mu\text{L}$ , ethanol 35.1 $\mu\text{L}$ , biocatalyst 108 mg, temperature 24.2 $^{\circ}\text{C}$ , hexane 2.05 mL, rotation – none;
<sup>c</sup> – prediction for the following reaction conditions: canola oil 206.1 μL, ethanol 45.6 μL, biocatalyst 62.5 mg, temperature 29.2 $^{\circ}$ C, hexane 1.81 mL, rotation – none; <sup>d</sup> – prediction for the following reaction conditions: canola oil 96.3 μL, ethanol 30 μL, biocatalyst 64 mg, temperature 32 $^{\circ}$ C, hexane 2.82 mL, rotation – none.

The coefficient of determination for the two-way interaction model was 0.7872; thus, the variability of performance was explained by the model at 78.72%. The value of the F-statistic was 5.286 for (21 and 30) degrees of freedom for the Fisher-Snedecor distribution. The test probability was 0. Therefore, at the significance level *α* = 0.05, the working hypothesis regarding the lack of significance of the multiple correlation coefficient was rejected in favor of the alternative hypothesis; thus, amount of canola oil and biocatalyst, temperature, and amount of solvent had a significant effect on the performance of FAEE biosynthesis.

**Table 5.**
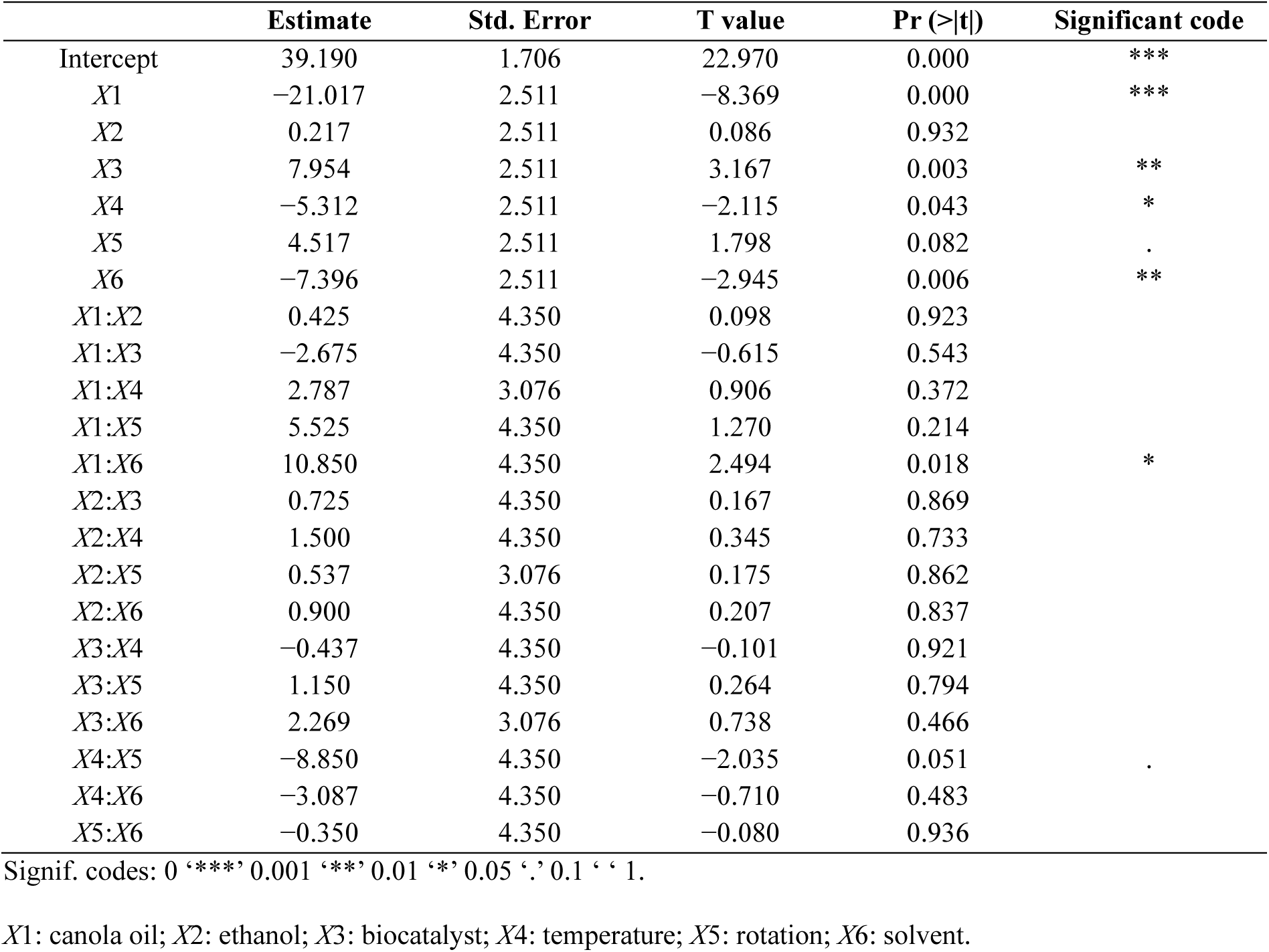
Structural parameter values of the TWI model, standard deviation, statistical value and test probability. Responses: FAEE yield (%)

|  | Estimate | Std. Error | T value | Pr (>\|t\|) | Significant code |
| --- | --- | --- | --- | --- | --- |
| Intercept | 39.190 | 1.706 | 22.970 | 0.000 | *** |
| X1 | -21.017 | 2.511 | -8.369 | 0.000 | *** |
| X2 | 0.217 | 2.511 | 0.086 | 0.932 |  |
| X3 | 7.954 | 2.511 | 3.167 | 0.003 | ** |
| X4 | -5.312 | 2.511 | -2.115 | 0.043 | * |
| X5 | 4.517 | 2.511 | 1.798 | 0.082 | . |
| X6 | -7.396 | 2.511 | -2.945 | 0.006 | ** |
| X1:X2 | 0.425 | 4.350 | 0.098 | 0.923 |  |
| X1:X3 | -2.675 | 4.350 | -0.615 | 0.543 |  |
| X1:X4 | 2.787 | 3.076 | 0.906 | 0.372 |  |
| X1:X5 | 5.525 | 4.350 | 1.270 | 0.214 |  |
| X1:X6 | 10.850 | 4.350 | 2.494 | 0.018 | * |
| X2:X3 | 0.725 | 4.350 | 0.167 | 0.869 |  |
| X2:X4 | 1.500 | 4.350 | 0.345 | 0.733 |  |
| X2:X5 | 0.537 | 3.076 | 0.175 | 0.862 |  |
| X2:X6 | 0.900 | 4.350 | 0.207 | 0.837 |  |
| X3:X4 | -0.437 | 4.350 | -0.101 | 0.921 |  |
| X3:X5 | 1.150 | 4.350 | 0.264 | 0.794 |  |
| X3:X6 | 2.269 | 3.076 | 0.738 | 0.466 |  |
| X4:X5 | -8.850 | 4.350 | -2.035 | 0.051 | . |
| X4:X6 | -3.087 | 4.350 | -0.710 | 0.483 |  |
| X5:X6 | -0.350 | 4.350 | -0.080 | 0.936 |  |

Signif. codes: 0 '\*\*\*' 0.001 '\*\*' 0.01 '\*' 0.05 '.' 0.1 ' ' 1.
X1: canola oil; X2: ethanol; X3: biocatalyst; X4: temperature; X5: rotation; X6: solvent.

The optimal conditions for biodiesel synthesis were used to determine the operational stability of the mycelium during subsequent catalytic cycles. The biocatalyst, in the form of mycelial *Penicillium* sp. 59, is characterized by high stability over 6-hour catalytic cycles. After six cycles, 94% of the initial activity was retained (Fig. 5).

**Figure 5.**
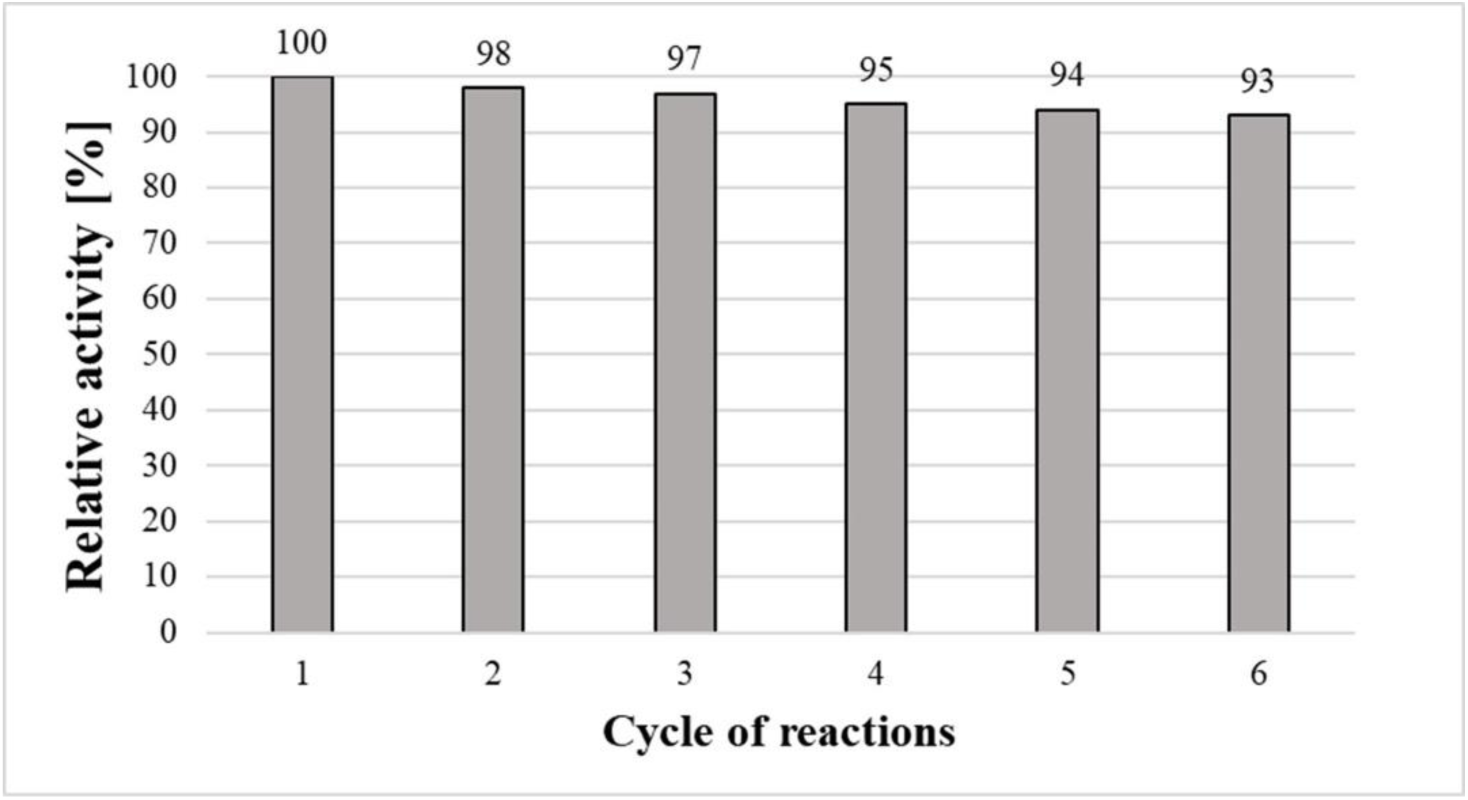
Reusability of *Penicillium* sp. 59 in a biodiesel synthesis.

## 4. DISCUSSION

Whole-cell biocatalysis has been successfully used in research on sustainable biodiesel production. The mycelium-bound lipases of mesophilic fungi have been of particular interest, such as those of *Rhizopus chinensis* [17], *Aspergillus niger* [9,10,18], *A*. *nomius*, *A*. *oryzae* [11], *R. oryzae* [19] and *Penicillium corylophilum* [20]. To date, mycelium-bound lipases from psychrotrophic fungi have not been used in research on the biocatalytic synthesis of biodiesel. The use of cold-adapted biocatalysts would make it possible to reduce the temperature of the process, thus increasing its profitability [13]. A psychrophilic lipase widely used in various biocatalytic processes is lipase B from *Candida antarctica* [21,22]. Its disadvantages include high price, need for immobilization, and heterologous expression in yeast cells for repeated use [23]. Therefore, research on new, efficient, and cheap psychrophilic biocatalysts for biodiesel synthesis is important. In this study, biocatalysts in the form of mycelium-bound lipases were screened from psychrotrophic fungi isolated from Arctic tundra soil and mesophilic fungi towards efficient biodiesel synthesis.

In the first stage of the study, the amount of biomass obtained, extra- and intracellular lipolytic activity, pH, amount of protein, specific extracellular activity, and synthetic activities of fatty acid methyl and ethyl esters were determined. The collection of a large amount of data was aimed at identifying variables that could be related to the synthetic activity of the mycelium. High extracellular lipolytic activity is not accompanied by high intracellular lipolytic or synthetic activity. In turn, low extracellular and intracellular lipolytic activity does not determine the low synthetic activity of mycelia in organic solvents. The low synthetic activity of fungal biocatalysts may be due to the high sensitivity of enzymes to organic solvents in which protein denaturation may occur [24]. Therefore, the screening of biocatalysts in the form of mycelia for esterification/transesterification reactions in an organic solvent cannot be based solely on agar plate tests or the determination of hydrolytic activity.

Fourier-transform infrared (FTIR) spectroscopy is a low-cost, simple, and fast technique that can be used to evaluate the synthetic activity of biodiesel biocatalysts in mycelial form. Analysis of the fungal biocatalyst spectra indicated the bands whose intensities could serve as predictors of biodiesel synthetic activity. The highest correlation, negative and positive, respectively was observed for two neighboring bands at 1745 cm^-1^ (*R* = −0.73) and 1699 cm^-1^ (*R*= 0.68). Similar correlations for these bands were noted during the immobilization of lipase from *Candida rugosa* on 3-(2,3-epoxypropoxy)propyltrimethoxysilane (EPOTS). The intensity of the bands in the amide I-III region was shown to decrease after immobilization compared to that of the free enzyme. The immobilized catalyst showed high hydrolytic activity towards *p*-nitrophenyl palmitate. This indicated favorable conformational changes in the protein [25]. In the present work, a high negative correlation was also noted for the intensity of the 1161 cm^-1^ band (*R* = −0.60). Lipase from *Burkholderia cepacia* immobilized by covalent binding with epichlorohydrin onto silica produced with a protonic ionic liquid showed a lower band intensity at 1161 cm^-1^ compared to the free enzyme. Carvalho et al. (2008) immobilized lipase from *Burkholderia cepacia* on two carriers, treated silica xerogels with protic ionic liquid and non-treated silica gel [26]. In the case of lipase immobilized on treated silica gel, a decrease in the intensity of the 1161 cm^-1^ band was observed. Moreover, biocatalysts with a lower intensity of this band were characterized by higher biodiesel synthetic activity compared to the biocatalyst immobilized on the untreated gel. Based on the above information, it can be concluded that an effective immobilization process is associated with a change in the FTIR spectra of the biocatalysts. The data collected in this study from various fungal species showed conservative changes within several bands. It should be noted that hundreds of data points are needed to fully correlate the intensity of individual bands with the synthetic activity of the fungi. However, we believe that this method may prove helpful for the rapid testing of large numbers of fungal strains.

A genetically modified biocatalyst, *Aspergillus oryzae*, in the form of whole cells that contained a gene encoding *Fusarium heterosporium* lipase, was used for the synthesis of FAME using soybean oil. The biodiesel yield was 94% after 96 hours of reaction at 30 °C [27]. Lipase from *Aspergillus oryzae* ST11 immobilized on a polyacrylonitrile nanofiber membrane in the form of magnetically cross-linked enzyme aggregates was characterized by efficient biodiesel synthesis [28,29]. The highest yield of FAME synthesis by immobilized lipase was 95% after 24 h at 37 °C. To date, a wild strain of *Aspergillus oryzae* in the form of whole cells has not been used as a biocatalyst for biodiesel synthesis. The wild-type strain of *A. oryzae* CBS 133.52 showed the same high synthetic activity (91.1% FAME and 94.5% FAEE yield) as the extracellular immobilized lipase from *Aspergillus oryzae* ST11. Moreover, the proposed process for obtaining the biocatalyst is much simpler. The efficiency of biodiesel synthesis using *R. oryzae* NRLL 3613 and *R. oryzae* DSM 63539 was similar to that obtained by other authors [19,30]. Our research has confirmed that the mesophilic strains of *Rhizopus oryzae* and *Aspergillus oryzae* are characterized by high biodiesel synthetic activity. The purpose of the use in this study of strains with known synthetic activity was to increase the amount of input data for the proper correlation of predictors of synthetic activity in strains. Among psychrotrophic strains, isolate 59 showed the highest efficiency. It showed a 57% yield of FAME and a 90% yield of FAEE synthesis. Due to the high activity of the unknown strain, we decided to carry out an analysis of phylogenetic affiliation and optimize the biocatalytic synthesis of fatty acid ethyl esters in further studies. The synthesis of FAEE has been optimized because of the easy acquisition of ethanol from agricultural sources, and FAEE contributes to lower exhaust gas emissions (including nitrogen oxides, CO_2_ and smoke density) than FAME [31].

Phylogenetic studies have shown that isolate 59 belongs to the *Penicillium* genus. The highest similarity of the examined sequence was with *Penicillium echinulatum* T3, 10-3 (accession number KP411588). However, this species is not recognized as a biocatalyst for biodiesel synthesis. Zhang et al. (2011) used lipase from *Penicillium expansum* to synthesize biodiesel in ionic liquids [32]. After 20 h of reaction with 8% biocatalyst content at 37 °C, an 86% yield of biodiesel synthesis was obtained. Huang et al. (2014), using 4% lipase from *Penicillium expansum* in an ionic liquid, obtained a 94% yield of biodiesel synthesis at 40 °C after 48 h of reaction [33]. Both processes require agitation of the system at 220 RPM. A biocatalyst in the form of whole cells of the genus *Penicillium* has not yet been used for the synthesis of biodiesel. However, the hydrolytic activities of these biocatalysts are known. Romero et al. (2014) investigated the activity and stability of a mycelium-bound lipase of *Penicillium corynophilum* [20]. The research shows that the biocatalyst in this form has a hydrolytic activity of 0.2 U/g of mycelium and shows high stability in organic solvents. Mycelium-bound lipase from *Penicillium citrinum* showed a high hydrolytic activity of 236 U/g and was used in the hydrolysis of vegetable oils [12]. The psychrotrophic strain *Penicillium* sp. 59 had a hydrolytic activity of 42 U/g of dry weight. However, as the results show in this work, based on hydrolytic activity, it cannot be unequivocally stated whether *P. corynophilum* and *P. citrinum* strains would also show synthetic activity.

*Penicillium* sp. 59 and *A. oryzae* as whole-cell biocatalysts showed the highest synthetic activities in highly hydrophobic solvents (hexane and heptane). Hexane was used as the solvent in the further optimization stage of biodiesel synthesis using *Penicillium* sp. 59. The use of ethyl acetate to initiate interesterification results in a decrease in the yield of the synthesis to approximately 25%. Importantly, the *Penicillium* sp. strain was characterized by a higher resistance to ethanol compared to *A. oryzae* (Fig. 4). Carrying out the reaction only in ethanol did not lead to a complete decrease in activity, as is the case with *A. oryzae*.

Optimization of biodiesel synthesis shortened the reaction time from 24 h to 6 h. At that time, the maximum real efficiency was 83.3%, with a projected efficiency of 85% from the TWI model. In each of the created models (FO, TWI, SO, and PQ), the factors that did not affect the efficiency of the synthesis were the volume of ethanol and the mixing of the system. The lack of ethanol effect probably resulted from the fact that the minimum dose (−1) of EtOH was sufficient for complete transesterification of the maximum dose (1) of used canola oil and from the resistance of the biocatalyst to ethanol at each dose. The TWI model best describes the biosynthesis of FAEE because of the similarity of the predicted value to the real value. Compared to other models, the TWI model is the most economical because the largest volume of oil is transesterified in one cycle, and it runs at a relatively low temperature of 24.2 °C. Moreover, the lack of effect of stirring the reaction mixture on efficiency reduces the energy necessary to power the stirrers. Notably, the overall cost of obtaining the biocatalyst is also low because of the low temperature of the culture.

An important factor that largely determines the profitability of the process on an industrial scale is the possibility of multiple uses of biocatalysts. The mycelium of Penicillium sp. 59 is characterized by high operational stability, and after six catalytic cycles, it loses only 7% of its initial activity. Equally high operational stability was demonstrated by whole-cell *Aspergillus* spp. RBD01 [29] and *A. oryzae* FH lipase [34], which were used for the synthesis of methyl esters.

## 5. CONCLUSION

Among the 29 tested strains, freeze-dried psychrotrophic mycelium of *Penicillium* sp. 59 showed high biocatalytic activity in the synthesis of fatty acid ethyl esters. The biocatalyst can be successfully used to design a cost-effective process on an industrial scale because of its high efficiency at room temperature, good stability in organic solvents, lack of need to mix, and possibility of repeated use. This research showed that it is impossible to unequivocally judge the synthetic activity of the biocatalyst based on its hydrolytic activity. Preliminary studies indicate that Fourier Infrared Spectroscopy (FTIR) may be a helpful technique for the rapid screening of fungi activity for biodiesel synthesis.

## Author Contributions

Conceptualization, M.K.; methodology, M.K., M.T., A.M-G.; software, M.K.; validation, M.K.; formal analysis, M.K.; investigation, N.J., W.J., S.G., E.P., M.K., K.G., A.S., A.Ś., L.C., A.B., A.M-G.; resources, M.K.; data curation, M.K.; writing—original draft preparation, M.K. N.J., W.J., S.G., E.P., K.G. L.C.; writing—review and editing, M.K., M.T.; visualization, M.K.; supervision, M.K., M.T.; project administration, M.K.; funding acquisition, M.K., M.T. All authors read and approved the manuscript.

## ACKNOWLEDGEMENTS

The authors would like to thank Maria Curie-Skłodowska University in Lublin, Poland for providing institutional funds to support this work.

